# Complex sphingolipids are essential for cell division and plasmodesmal development in the moss *Physcomitrium patens*

**DOI:** 10.1101/2024.11.01.621568

**Authors:** Linus Wegner, Cornelia Herrfurth, Ivo Feussner, Katrin Ehlers, Tegan M. Haslam

## Abstract

Developmental patterning and organ structure are elegantly simple in the moss *Physcomitrium patens*. In molecular genetic studies, this facilitates both the cultivation of severe mutant alleles and their phenotypic characterization. Essential membrane lipids, such as complex phosphosphingolipids (in plants, glycosyl inositol phosphorylceramides, GIPCs), have been difficult to functionally characterize due to non-viable and pleiotropic phenotypes of mutants affected in their synthesis in *Arabidopsis thaliana*. Following the isolation and biochemical characterization of mutants affected in GIPC synthesis in *P. patens*, including *sphinganine-C4-hydroxylase* (*s4h*/*sbh*) and *inositol phosphorylceramide synthase* (*ipcs*), we now report some of their morphological, histological, and cytological phenotypes. We observed alteration in cell division, expansion, and differentiation. Specifically, the *s4h* knock-out mutant had abnormal cell division planes, as well as irregular depositions attached to cell walls. Severe *ipcs* mutant alleles showed frequent incomplete cell divisions, causing compromised cell autonomy as demonstrated by intercellular motility assays. These phenotypes suggest that sphingolipids impact both the orientation and proper formation of the cell plate during cytokinesis. Transmission electron microscopy revealed dramatic plasmodesmal structural defects in all three mutants, however, qualitative aspects of plasmodesmal transport do not seem to be severely impacted. Our methods can be used as a toolkit for quantifying growth, and specifically cell division and plasmodesmal phenotypes in mosses; our present results elucidate the specific contributions of GIPCs to fundamental cell functions. Finally, the severity of the observed defects in cell functions and ultrastructure highlight the resilience and utility of *P. patens* for studying basic cellular functions and severe mutant phenotypes.

## Introduction

Glycosyl inositol phosphorylceramides (GIPCs) are complex phosphosphingolipids that are essential and abundant components of plant cell membranes (Tjellström et al., 2010; Cacas et al., 2016; Bahammou et al., 2023) (Figure 1A, B). Genetic manipulation of different plant species, including *Arabidopsis thaliana*, *Oryza sativa*, and *Medicago truncatula*, has produced mutants with modified GIPC content that exhibit abnormalities in their interactions with symbiotic microorganisms (Moore et al., 2021), pathogen resistance (Lenarčič et al., 2017), and responses to abiotic environmental conditions (Jiang et al., 2019). GIPC-deficient mutants also usually show obvious growth defects in the absence of environmental stresses, and knock-out lesions that block GIPC assembly entirely cannot be recovered (Wang et al., 2008; Rennie et al., 2014; Ito et al., 2021). This suggests that in addition to contributing to plant interactions with their environment, GIPCs contribute to normal physiology and development, and underscores the notion that they are essential.

**Figure 1:**
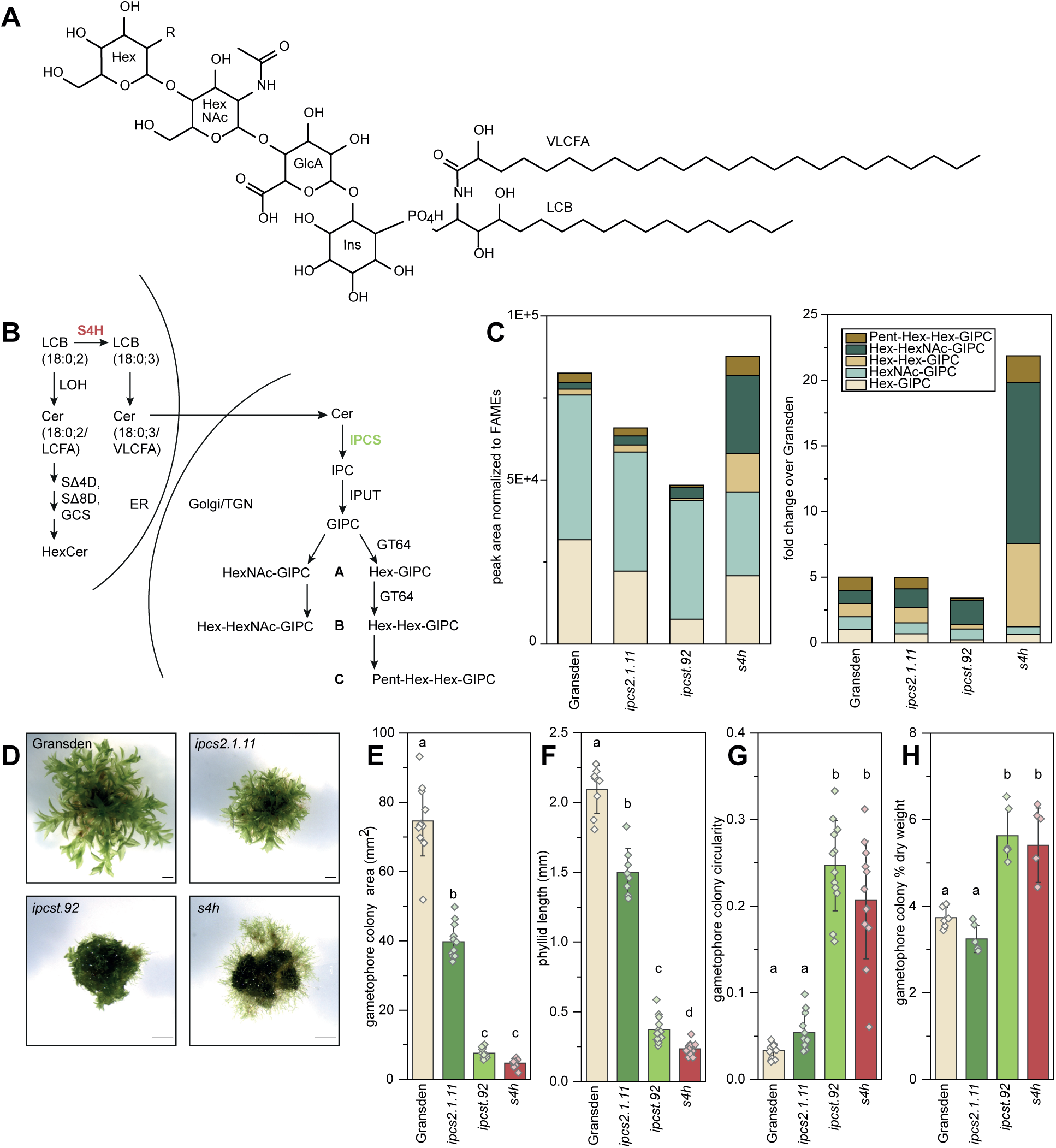
Altered sphingolipid content of mutant gametophores affects their overall growth. (**A**) Chemical structure of a sphingolipid, specifically Hex-HexNAc-GIPC, 18:0;3/24:0;1, where 18:0;3 represents the long-chain base (LCB) with 18 carbons in length, no unsaturated bonds, and three hydroxylations, and 24:0;1 the very-long-chain fatty acid (VLCFA) moiety with 24 carbons in length, no unsaturated bonds, and a single hydroxylation. (**B**) Simplified schematic of GIPC biosynthesis. Ceramide (Cer) assembly and backbone modifications occur on the endoplasmic reticulum, including SPHINGANINE-4-HYDROXYLASE (S4H, pink) hydroxylation of 18:0;2 LCBs to 18:0;3. In wild-type *P. patens*, only 18:0;3 LCBs are incorporated in GIPCs. Ceramides containing an 18:0;3 LCB and a VLCFA move into the Golgi and *trans*-Golgi network (TGN) where phospho-sugar headgroup additions build GIPCs. INOSITOL PHOSPHORYLCERAMIDE SYNTHASEs (IPCSs, green) catalyze the committed step in GIPC assembly. (**C**) GIPC profiles of the mutants, based on ultra-high pressure liquid chromatography coupled with nanoelectrospray and tandem mass spectrometry (UPLC-nanoESI-MS/MS) quantification of lipids extracted from microsome enrichments from gametophores. Different GIPC class amounts are sums of individual MRM peak areas, normalized to the fatty acid content in each sample determined by FAMEs analysis (left). On the right, the same dataset, but processed as fold-change compared to the amount in wild type, where the amount of each GIPC type is set to a value of 1. The complete dataset is presented in Supplementary Figure S1.(**D**) Gametophores of the wild-type Gransden, *ipcs2.1.11* single mutant, *ipcst.92* triple mutant, and *s4h* mutant, shown at different magnifications. All scale bars 1 mm. (**E**) Gametophore colony area, (**F**) Phyllid length, and (**G**) gametophore colony circularity after approximately two months of growth. (**H**) % dry weight of gametophore tissue produced for lipid analysis. Bars are averages of 12 (**E**&**G**), 8-14 (**F**), and 6 (**H**) replicates with standard deviation, letters indicate significance at p < .05 determined by one-way ANOVA with Tukey’s *post-hoc* test. LOH, LAG ONE HOMOLOG; SΔ4D, SPHINGOLIPID DELTA 4 DESATURASE; SΔ8D, SPHINGOLIPID DELTA 8 DESATURASE; GCS, GLYCOSYL CERAMIDE SYNTHASE; IPUT, INOSITOL PHOSPHORYLCERAMIDE GLUCURONOSYLTRANSFERASE; GT64, GLYCOSYL TRANSFERASE 64.

Defining precise roles of GIPCs in growth and development is challenging for two reasons. First, most studies on GIPC functions have been conducted on vascular plant models. Their complex organ structures and developmental patterns can be difficult to analyze, and mutants can be non-viable when even a single developmental step is blocked by a given mutation. Second, GIPCs are synthesized from ceramides, the defining lipidic backbone common to sphingolipids. Therefore, blocking steps in headgroup assembly not only results in reduced GIPC levels, but also increased levels of free ceramides. As ceramides are potent signals that trigger programmed cell death (Berkey et al., 2012; König et al., 2022), it can be difficult to resolve which mutant phenotypes are caused by substrate excess and which by depleted product.

GIPCs accumulate together with sterols into lipid-ordered microdomains within the plasma membrane, which can direct the organization of other protein and lipid membrane components (Gronnier et al., 2016). Specifically, recent work on artificial microRNA (amiRNA) *IPCS* transgenic lines of *A. thaliana* suggested that GIPCs drive the localized depletion of the phosphoinositide phosphoinositol-4-phosphate PI(4)P (Ito et al., 2021). The development and application of fluorescent probes with affinity for specific phosphoinositides has revealed myriad impacts of their specific localizations on fundamental cell biology (Simon et al., 2014; Noack and Jaillais, 2017; Markovic and Jaillais, 2022). Therefore, by mediating PI(4)P distribution, GIPCs could be expected to have an impact on these processes as well. Further, the very-long-chain fatty acid (VLCFA) moieties of GIPCs have been shown to be essential for normal endomembrane trafficking (Molino et al., 2014; Wattelet-Boyer et al., 2016), likely through a direct role in stabilizing highly-curved membranes that form during vesicle fusion (Schneiter et al., 2004; Molino et al., 2014).

Additionally, several studies have suggested that GIPCs are involved in the formation and function of plasmodesmal cell connections in plants (Zhang et al., 2022). Enrichment of plasmodesma-derived plasma membrane from cell walls of *A. thaliana* suspension cell cultures revealed increased GIPC content relative to the total plasma membrane (Grison et al., 2015). Further, mutants of *A. thaliana PHLOEM LOADING MODULATOR* (*PLM*) displayed defects in plasmodesmal development and intercellular connectivity, as well as a mild reduction in GIPC levels (Yan et al., 2019). PLM shares sequence identity and conserved motifs with *A. thaliana* INOSITOL PHOSPHORYLCERAMIDE SYNTHASEs (IPCSs), which catalyze the committed step in GIPC synthesis. Therefore, although the biochemical activity of PLM remains uncertain, its sequence and mutant phenotype suggest it may contribute to GIPC accumulation in a cell type- or membrane domain-specific manner.

*Physcomitrium patens* is a good genetic system for isolating and studying strong mutant phenotypes that severely affect growth and development. There are routine lab protocols for not only propagating *P. patens* tissue vegetatively, but also carrying out precise genome manipulation without going through a reproductive cycle (Schaefer, 2001; Lopez-Obando et al., 2016; Collonnier et al., 2017); as such it is possible to generate and cultivate mutants with severe developmental defects. Additionally, the filamentous protonemal growth and single-cell thick phyllids of moss gametophores are easy to image and to monitor for phenotypes related to cell division, differentiation, and growth.

Profiling *P. patens* sphingolipids with targeted lipidomics revealed the presence of GIPCs containing hexose sugars (Hex) as well as *N*-acetylated hexoses (HexNAc), within the same tissues (Resemann et al., 2021; Steinberger et al., 2021). Additionally, A-, B-, and C-series GIPCs could be detected having 1, 2, or 3 sugars (Hex, HexNAc, or pentose [Pent]), respectively, linked to the glucuronic acid moiety of the headgroup (Steinberger et al., 2021) (See structure, Figure 1A, and simplified pathway, Figure 1B). We previously identified several GIPC-deficient *P. patens* mutants by targeting the *IPCS* gene family, which consists of three members in the moss (Haslam et al., 2024). We were especially interested in an *ipcs2* single mutant that showed a moderate decrease in Hex-GIPCs (*ipcs2.1.11*), and an *ipcs1 ipcs2 ipcs3* triple mutant that showed severe reductions in Hex-GIPCs and Hex-Hex-GIPCs (*ipcst.92*). While these alleles caused relatively strong decreases in A- and B-series GIPCs with hexose headgroups, they showed only modest reductions in HexNAc-containing GIPCs. The *ipcst.92* mutant was also unusual in that it showed no significant increase in ceramide substrate despite the GIPC product depletion, and an exponential accumulation of the direct product of IPCS activity, inositol phosphorylceramides. In our previous work, we proposed a model in which one of the three mutagenized *ipcs* loci in this triple mutant is still expressed and active, but produces and accumulates the inositol phosphorylceramide product in a way or in a location where it is inaccessible for further glycosylations to produce mature GIPCs (Haslam et al., 2024). This would explain both the inositol phosphorylceramide accumulation and the wild type-like levels of ceramide in the mutant. We also previously generated *sphinganine-C4-hydroxylase* (*s4h*) mutants (Gömann et al., 2021) (also known as *sphingoid base hydroxylase*, *sbh* (Steinberger et al., 2021)). S4H hydroxylates di-hydroxy (18:0;2 or d18:0) long-chain bases (LCBs) to tri-hydroxy LCBs (18:0;3 or t18:0), which are preferentially used in the synthesis of GIPCs. Perhaps surprisingly, among the different changes in the sphingolipid profile of *s4h* mutant, we noticed a 5- to 6-fold increase in B-series GIPCs, including Hex-Hex-GIPCs and Hex-HexNAc-GIPCs, while the A-series GIPCs were not significantly altered compared to the wild type.

Together, this selection of moss mutants presents a good toolkit to investigate the impacts of different GIPC defects on growth and development, in a model system that is both easy to image and particularly tolerant of basic physiological and developmental deficiencies. We quantified macroscopic growth of gametophores, cell division and differentiation in the leaf-like phyllids, and ultrastructural differences between the wild type and mutant cells. We observed strong phenotypes related to cell division in both *ipcst.92* and *s4h*, and malformation of plasmodesmata in all mutant alleles. The physiological consequences of these defects were further investigated using intercellular motility assays with protein and small chemical fluorescent probes. Our results identify roles of GIPCs in fundamental plant cell biology, in a bryophyte model that provides a complementary and phylodiverse comparison to *A. thaliana*.

## Results

### *ipcs* and *s4h* mutants cause substantial and distinct modifications to the GIPC profile of *P. patens*

We first confirmed the metabolic deficiencies of our mutants of interest, after cultivating and processing material from all genotypes in parallel, under uniform growth conditions. This supported our previously-reported analyses (Gömann et al., 2021; Haslam et al., 2024), with several substantial and significant differences to GIPC profiles of the mutants motivating our interest (Figure 1C). There are also minor discrepancies in the present work compared to our previous reports; this should be due at least in part to different growth conditions, and to the fact that we used mature gametophores for analysis, whereas protonema was used in (Gömann et al., 2021).

Absolute quantification of GIPCs is not possible in the absence of standards. Therefore, we expressed differences in GIPC amounts between samples as MRM peak areas normalized against a total FAMEs measurement. This provides a rough proxy for the total amount of acyl lipid per sample (Figure 1C, Supplementary Figure S1). Additionally, we expressed GIPC amounts in fold-change relative to the wild-type measurement for that lipid class, with the wild-type value for each class of GIPC arbitrarily set to one; this provides a different perspective of the relative shift in each class (Figure 1C). Although neither quantification is absolute, in the absence of standards, this is the best and most objective analysis we can devise.

We confirmed the mild reductions in Hex-GIPCs of the *ipcs2.1.11* mutant, which were decreased approximately 30 % relative to wild type (Supplementary Figure S1A), and resulted in an approximately 20 % reduction of the overall GIPC levels (Figure 1C). While not statistically significant in this measurement, this difference was reproducible between multiple experimental replicates. In the more severely-impacted *ipcst.92* triple mutant, we observed an approximately 70 % reduction in Hex-GIPCs (Supplementary Figure S1A), and similar reductions in Hex-Hex-GIPCs (Supplementary Figure S1B) and Pent-Hex-Hex-GIPCs (Supplementary Figure S1C), compared to wild type, while levels of HexNAc-decorated GIPCs were unchanged (Supplementary Figure S1D, E). Overall, this resulted in an approximate 40 % reduction of total GIPCs in this mutant (Figure 1C). The conspicuous trend of *ipcs* lesions having little or no impact on HexNAc-containing GIPCs suggests that the synthesis of these could be regulated differently than for GIPCs decorated with only hexose(s) and pentose. We also confirmed the puzzling GIPC chemotype of the *s4h* mutant, namely the dramatic increase in B-series GIPCs (6-fold increase in Hex-Hex-GIPCs, 12-fold increase in Hex-HexNAc-GIPCs, Figure 1C, Supplementary Figure S1B, E), and further observed a more moderate increase in C-series GIPCs (2-fold increase in Pent-Hex-Hex-GIPCs, Figure 1C, Supplementary Figure S1C). We also observed mild, though insignificant, reductions in A-series GIPCs in *s4h* (Figure 1C, Supplementary Figure S1A, D); altogether these changes resulted in a slight increase in total GIPC levels in *s4h* relative to wild type (Figure 1C).

Importantly, the metabolic phenotypes of *ipcs* and *s4h* mutants are not limited to GIPCs. As we previously reported, *ipcst.92* strongly accumulates IPCs (Supplementary Figure S1F). However, a transgenic line we generated in the wild-type background (a partial genetic replica of *ipcst.92,* “wt>t92”), using only one of the three mutagenized loci from *ipcst.92* (*ipcs2*), maintained a wild type-like growth phenotype while producing the strongly increased IPC levels characteristic of *ipcst.92*. That this line lacks all developmental phenotypes of *ipcst.92* indicates that it is the cumulative loss of IPCS activity in the *ipcst.*92 triple mutant, not IPC accumulation, that correlates to its growth deficiencies (Haslam et al., 2024). We also observed no substantial difference in the free ceramide and HexCer levels of the *ipcs* mutants (Supplementary Figure S1G-I).

The metabolic phenotype of the *s4h* mutant is especially complex in its impact on different sphingolipid classes. We observed near-complete depletion of all sphingolipids containing an 18:0;3 LCB moiety, which are replaced with 18:0;2 LCB moieties (Supplementary Figure S1A-F). In the case of free ceramides, this compensation was incomplete, with free ceramides accumulating to approximately 20 % of the wild-type level (Supplementary Figure S1G). Additionally, HexCer levels in *s4h* nearly doubled compared to wild type (Supplementary Figure S1H), and strikingly, contain a more diverse fatty acid profile (Supplementary Figure S1I). In summary, all three mutants have strong GIPC-related phenotypes, with *ipcs* mutants showing overall GIPC reduction, and differences in Hex- vs. HexNAc-GIPCs. The *s4h* mutant displays a strong shift to more highly-glycosylated GIPCs. The sphingolipid phenotypes of the *ipcs* mutants are essentially limited to GIPCs, whereas the phenotype of *s4h* is complex, and its growth and development may be impacted by other changes in its sphingolipid profile.

### The *ipcs2.1.11*, *ipcst.92*, and *s4h* mutants have obvious and distinct morphological phenotypes

All three mutants have obvious morphological phenotypes (Figure 1D). These were quantified based on images of 2-month-old gametophores of different genotypes grown together on the same plates. Gametophore colony area is significantly reduced in all three genotypes, with a near 50 % reduction in *ipcs2.1.11*, and approximately 90 % reductions in both *ipcst.92* and *s4h* (Figure 1E). *ipcs2.1.11* overall appears morphologically similar to wild type, but smaller. In contrast, *ipcst.92* and *s4h* are both dwarf and morphologically unusual. Gametophore expansion was reduced in *ipcst.92* and *s4h* (Figure 1E), to a degree that was more severe than reduction of individual phyllid size (Figure 1F). This discrepancy between colony size reduction and phyllid length reduction gave these two mutants a dense, round appearance compared to the branched and stellate wild type and *ipcs2.1.11*. The altered form of *ipcst.92* and *s4h* can be quantified as circularity, which increased nearly 10-fold in these two mutants compared to wild type (Figure 1G). Additionally, the % dry mass of *ipcst.92* and *s4h* was significantly increased relative to wild type, by approximately 30 % (Figure 1H), presumably due at least in part to the relative increase in apoplastic vs. symplastic material that would occur with reduced cell size, and corresponding increased surface area-to-volume ratio.

Comparing *ipcst.92* and *s4h* to each other, there are also clear differences. Though both mutants required more time to undergo the transition from protonema and gametophore growth, thereafter, *ipcst.92* grew uniformly as such. In contrast, the *s4h* mutant never established homogeneous gametophore growth, and the gametophores that were initiated were morphologically irregular.

### Cell division, expansion, and differentiation are all altered compared to wild type in mutant phyllids

Phyllid histological features and differentiation into distinct cell types (Dennis et al., 2019; Lin et al., 2021) were next observed in surface view (Figure 2A-G), and in crystal violet-stained semi-thin sections of the median regions (Figure 2H-U). There are several distinct cell types in phyllids. First, phyllid margin cells (also called edge cells), have an elongated form and protrude slightly at the apical end, producing a rounded serration for the margin. Second, stereid and hydroid cells in the hadrom (a “midrib” in analogy to vascular plants), are elongated, straight, and stereids have thicker cell walls than the other cell types. The remaining cells between the hadrom and margin are considered here simply as parenchyma, and vary substantially in size, but generally have a shorter aspect ratio than the margin, stereid, and hydroid cells, and are more densely filled with chloroplasts. In some literature, parenchyma cells are further sub-divided into base and tip cells, and are distinguished based on their different shape and size (Dennis et al., 2019; Lin et al., 2021).

**Figure 2:**
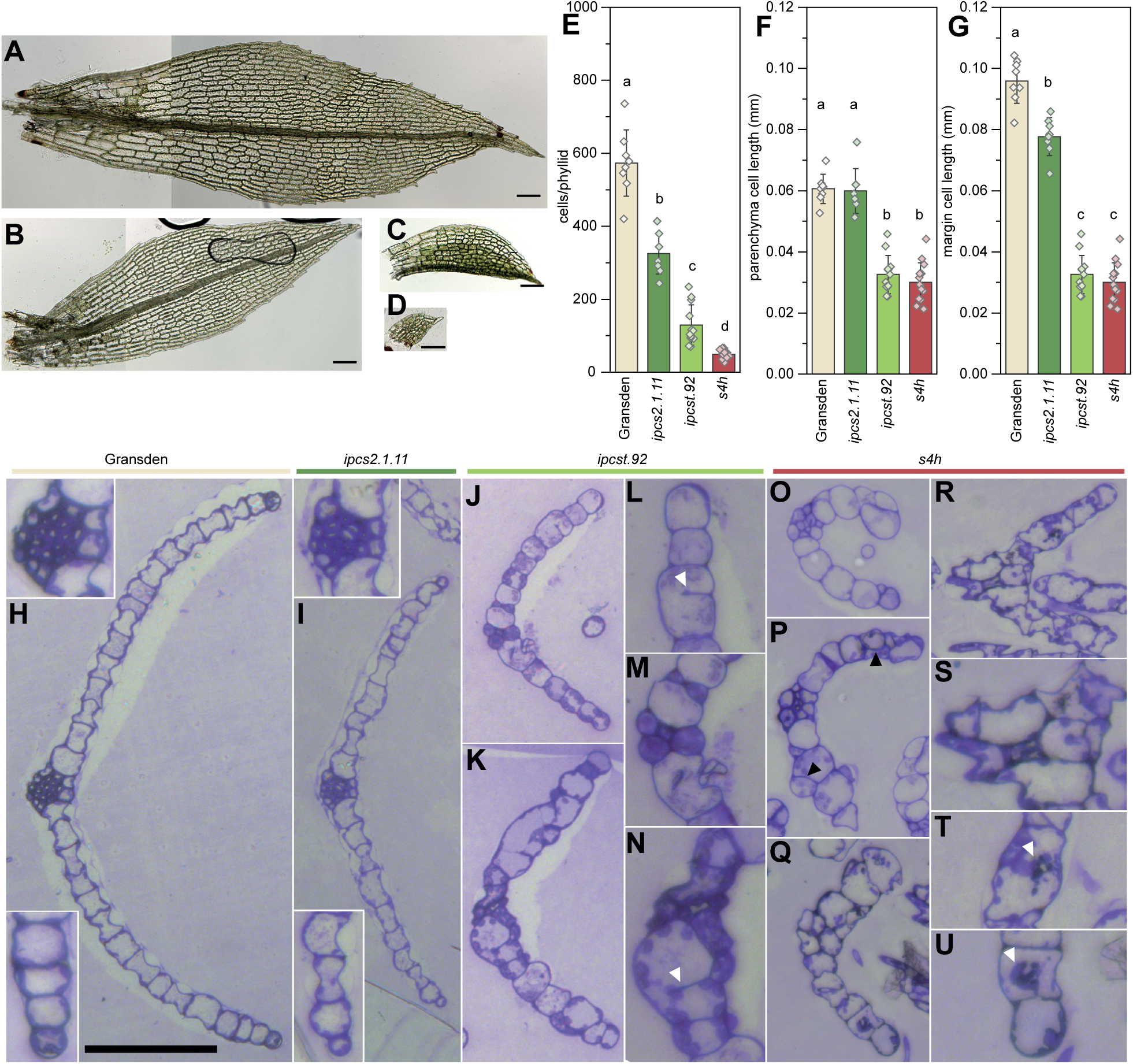
Representative images of wild-type Gransden (**A**), *ipcs2.1.11* (**B**), *ipcst.92* (**C**), and *s4h* (**D**) used for phyllid and cell measurements (**E-G**). All scale bars represent 0.1 mm. Images are stitched from multiple snapshots taken of different phyllid regions, and in different focal planes. (**E**) Number of cells per phyllid. (**F**) Length of phyllid parenchyma cells. (**G**) Length of phyllid margin cells. Bars are averages of 8-14 phyllids with standard deviation, letters indicate significance at p<.05 determined by one-way ANOVA with Tukey‘s *post-hoc* test. Wild-type Gransden (**H**), *ipcs2.1.11* single mutant (**I**), *ipcst.92* triple mutant (**J-N**), *s4h* mutant young (**O,P**) and mature phyllids (**Q-U**). The scale bar corresponds to 100 µm in the overviews (**H, I, J, K, O-R**); and to 50 µm in the insets of (**H**, **I**), and in (**L-N**, **S-U**), which show enlarged details of the hadroms and the phyllid margin cells, as well as mutant peculiarities. Incomplete cell walls are marked with white arrowheads in (**L**, **N**), oblique and periclinal divisions with black arrowheads in (**P**), and irregularly shaped, densely stained material is marked with white arrowheads in (**T**, **U**).

Dramatic reductions were clear in phyllid cell numbers for all mutants, with *ipcs2.1.11* reduced to 55 %, *ipcst.92* to 20 %, and *s4h* to 10 % of the cell number per phyllid compared to the wild type (Figure 2E). Average cell length was calculated by measuring individual phyllids and dividing the length by the number of cells in a column spanning the length of the phyllid (Figure 2F, G). We could clearly distinguish between parenchyma and margin cells in the Gransden wild type and in *ipcs2.1.11*, and so these cell types could be measured separately. In *ipcst.92* and *s4h*, there was no consistent, distinct shape to cells at the phyllid margin; additionally, cell columns within the phyllid were difficult to track. Therefore, only an edge row of cells was used for quantification of these two genotypes. By comparison to wild-type parenchyma or margin cells, *ipcst.92* and *s4h* phyllid cells were reduced to between 30-50 % of the normal length (Figure 2F, G). For *ipcs2.1.11*, only margin cells were approximately 15 % shorter (Figure 2G), whereas parenchyma showed no significant difference (Figure 2F). Overall, the reduction in phyllid size was primarily due to reduced cell number for *ipcs2.1.11*, whereas for *ipcst.92* and *s4h*, both cell length and cell number are substantially reduced.

In median sections through wild-type Gransden phyllids (Figure 2H), the water-conducting hydroids and the thick-walled stereids located in the hadrom (upper inset in Figure 2H) separated the two halves of the phyllid, which consisted of parenchyma, and margin cells at the outer edge (lower inset in Figure 2H), in a single cell layer. Anticlinal cell walls were regularly-spaced and occurred at near-perfect right angles to the phyllid surface. The phyllid anatomy and histology of the *ipcs2.1.11* single mutant (Figure 2I) did not differ considerably from the wild type, with an easily discernible hadrom (upper inset in Figure 2I), differentiated margin cells, and normally-shaped but fewer parenchyma cells (lower inset in Figure 2I).

Phyllid anatomy was severely affected in the *ipcst.92* triple mutant, with obvious developmental defects in the hadroms, which were composed of fewer, but often wider cells (Figure 2J, K, M). Only few, if any, cells developed the characteristically thick stereid cell walls. Apart from the reduced parenchyma cell numbers and lengths (Figure 2E-G, J, K), further observation of *ipcst.92* phyllid sections hinted at a general defect in the control of cell division: we often observed strikingly wide cells with incomplete anticlinal walls, which extended from one side of the parental cell wall and ended bluntly in the cell, thereby failing to separate adjacent cells completely from each other (Figure 2L, N).

Phyllids of the *s4h* mutant also showed severe histological defects, with either poorly- or indiscernible hadroms, and wide variation in parenchyma cell width and shape throughout the phyllid cross sections (Figure 2O-S). They had striking oblique and periclinal division planes, leading to partially-multilayered phyllids. Another characteristic feature of the *s4h* mutant phyllid cells was irregularly shaped, tufted aggregates of material, darkly stained with crystal violet, whose direct attachment to the cell walls could be seen within serial sections (Figure 2T, U).

Reduced size of the mutant phyllids and reduced cell number per phyllid (Figure 2A-E) could be easily correlated to reduced numbers of cell columns per phyllid in these cross sections (Figure 2H-U), approximately 15 per phyllid half in Gransden, 10 in *ipcs2.1.11*, 8 in *ipcst.92*, and 6 in *s4h*. Perhaps surprisingly, the reductions in average cell length measured in whole phyllids (Figure 2F, G) did not correlate with widths observed in semi-thin sections (Figure 2H-U). We infer that whatever factors caused stunted cell elongation in *ipcst.92* and *s4h* did not substantially impact lateral cell expansion.

### TEM investigations reveal additional unique phenotypes in *ipcs* and *s4h* mutants

We sought to better understand the unusual phenotypes observed via light microscopy by TEM. Observation of the blunt-ended anticlinal cell walls in the *ipcst.92* mutant (Figure 3A, B) revealed the presence of plasmodesmata (black arrowhead in Figure 3A), as well as regular internal layering and the presence of a middle lamella (white arrowhead in Figure 3B). Structurally, they were not obviously different from other cell walls in the *ipcst.92* mutant, or walls of wild-type phyllids. While this phenotype requires further investigation, our observations of incomplete cell walls in *ipcst.92* suggest that this mutant fails to organize cell plate attachment to the parental cell wall to complete cytokinesis.

**Figure 3:**
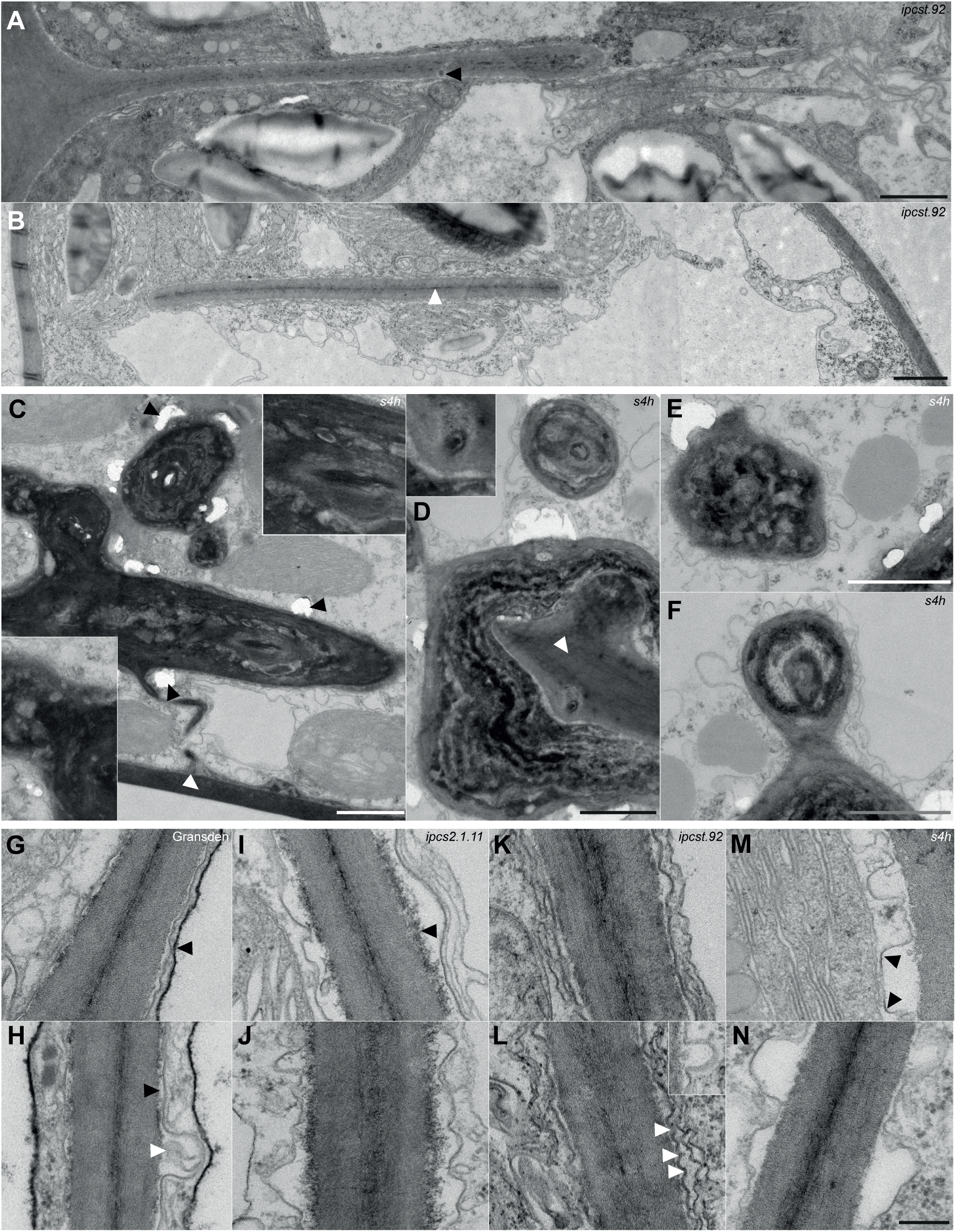
Ultrastructural peculiarities observed in the mutants *via* transmission electron microscopy. (**A**,**B**) TE micrographs showing incomplete cell walls, which end bluntly in mature phyllid cells of *ipcst.92*, and exhibit a regular layering of the middle lamellae (white arrowhead in **B**) and the subsequent cell wall layers as well as plasmodesmata (black arrowhead in **A**). Analyses of serial light-microscopy images revealed that apparently non-attached incomplete walls are connected to the outer cell wall in other planes of section. (**C**-**F**) TE micrographs of cytosolic regions of *s4h* mutant cells with irregularly-shaped, tufted aggregates. Attachment to the outer cell walls can be seen in some sections (white arrowheads in **C** and **D**), which included plasmodesmata (**D**, inset). The appositions usually had a layered substructure composed of both light and dark materials (**C**, insets). They likely had a hard consistency, since ultrathin sections often cracked in the vicinity of the aggregates (black arrowheads in **C**). All scale bars in (**A-F**) represent 1 µm, insets have a two-fold magnification. (**G**-**N**) Representative TE micrographs showing peculiarities of the attachment of the plasma membrane to the cell wall. In Gransden (**G**,**H**) the plasma membrane lines the cell wall closely (black arrowhead in **H**), and single vesicles were regularly found to fuse with the plasma membrane (white arrowhead in **H**). The tonoplast membrane appeared unusual in that it was more electron-dense (black arrowhead in **G**) than is typically observed in wild type *P. patens* (Reute), and in the mutants from this sample set. In *ipcs2.1.11* (**I**,**J**), the plasma membrane had a wavy shape and was detached from the cell wall over wide areas. The cell wall appears unusual in that its youngest layers were irregularly fringed (black arrowhead in **I**). The plasma membrane of *ipcst.92* (**K**,**L**) was also wavy and detached from the cell wall over large areas, with more frequent but smaller membrane bends (white arrowheads in **L**). The inset in (**L**) shows the fusion of a small vesicle with the plasma membrane. The shape of the plasma membrane of the *s4h* mutant (**M**,**N**) resembles that of the *ipcs2.1.11* single mutant, but detached membrane areas were more common, more deeply invaginated, and often included particularly straight and angular membrane sections with higher electron density (black arrowheads in **M**). Scale bar for (**G**-**N**), shown in (**N**), is 200 nm.

The intracellular aggregations in *s4h* were further investigated via TEM: They appeared irregular in form, mostly electron-dense, and had an intricate, layered sub-structure including some electron-lucent regions (Figure 3C-F). The ultrathin sections were often cracked directly adjacent to these aggregates (black arrowheads in Figure 3C), suggesting that they may have an especially hard consistency. They were often observed in contact with cell walls (white arrowheads in Figure 3C, D). The overall form of these depositions was reminiscent of aniline blue-stained structures previously observed surrounding cross walls in protonemal filaments of *s4h*, inferred to be abnormal callose depositions (Gömann et al., 2021; Ye and Zhong, 2022), as well as red-pigmented intracellular aggregates previously observed in phyllids of *s4h* (*sbh*) (Steinberger et al., 2021). The presence of both of these accumulations was confirmed in gametophores grown under our experimental conditions via light and fluorescent microscopy. Notably, aniline blue-stained and red-pigmented structures did not co-localize (Supplementary Figure S2).

Due to the chemical impact of GIPCs on plasma membrane-cell wall adherence (Voxeur and Fry, 2014), we attempted to visually observe contact between these structures in TE micrographs of our GIPC-deficient mutants (Figure 3G-N). In wild type, (Figure 3G, H), there were regions with close, parallel alignment between the plasma membrane and cell wall (black arrowhead in 3H), as well as vesicle contact sites where the plasma membrane is invaginated (white arrowhead in 3H). The tonoplast membrane seemed to be associated with granular vacuolar material and appeared unusually thick and darkly-stained (black arrowhead in 3G), even by comparison to similarly-treated *P. patens* Reute phyllids (Wegner and Ehlers, 2024). Tonoplast membranes with similar electron-density were only occasionally observed in *ipcs2.1.11.* In all three GIPC mutants, plasma membrane detachment from the cell wall could easily be observed, but with distinct characteristics in each mutant (Figure 3I-N). In *ipcs2.1.11* (Figure 3I, J), detachment occurred over large regions, and the plasma membrane had a wavy appearance. The inner layers of the cell wall often had an electron-dense fringe (black arrowhead in 3I). In *ipcst.92* (Figure 3K, L) large expanses of wavy, detached plasma membrane were obvious (white arrowheads in 3L); here the waves had a markedly shorter length, producing a densely rippled appearance, and the invaginations were not as deep as in the other two mutants. Plasma membrane detachments in *s4h* were also frequent (Figure 3M, N), and remarkably, the membrane curvature often was sharp to the point that it appeared angular, and the invaginations were deepest here among the three mutant genotypes. These angular membrane sectors appeared to have higher electron density within the otherwise weakly contrasted plasma membrane (black arrowheads in 3M).

### Plasmodesmal ultrastructure is impacted by modification of GIPC content

TE micrographs were also examined for alterations of plasmodesmal structures of the mutants. Although median phyllid sections were compared between genotypes, the strong developmental phenotypes of the *ipcst.92* and *s4h* mutants introduced some complexity to these comparisons given that plasmodesmal structure and function is known to progress alongside cell maturation. The specifics of this process were recently described for *P. patens* Reute (Wegner and Ehlers, 2024). Similar to angiosperms (Nicolas et al., 2017), young, narrow type I plasmodesmata typically have a uniform, approximately 25 nm diameter over their length, and internal structures cannot be discriminated. These plasmodesmata have a relatively high size exclusion limit (SEL) of at least 27 kDa, which allows for the movement of macromolecules between cells. In contrast, in older type II(-like) plasmodesmata characteristic of mature cells, internal substructures can clearly be seen, including the central ER-derived desmotubule and the cytosolic sleeve between the plasma membrane and desmotubule. These type II-like plasmodesmata are wider at approximately 45 nm in the dilated region located in the median cell wall layers (white arrows in Figure 4A-N), and show constricted ‘necks’ only at the orifices (white arrowheads in Figure 4A-N), where the plasma membrane and desmotubule membrane (black arrowheads in Figure 4A-N) are still locally difficult to distinguish. Type II plasmodesmata have a much lower size exclusion limit, approximately 1 kDa in *A. thaliana*, and therefore restrict the intercellular transport between older cells to small molecules (Nicolas et al., 2017; Wegner and Ehlers, 2024).

**Figure 4:**
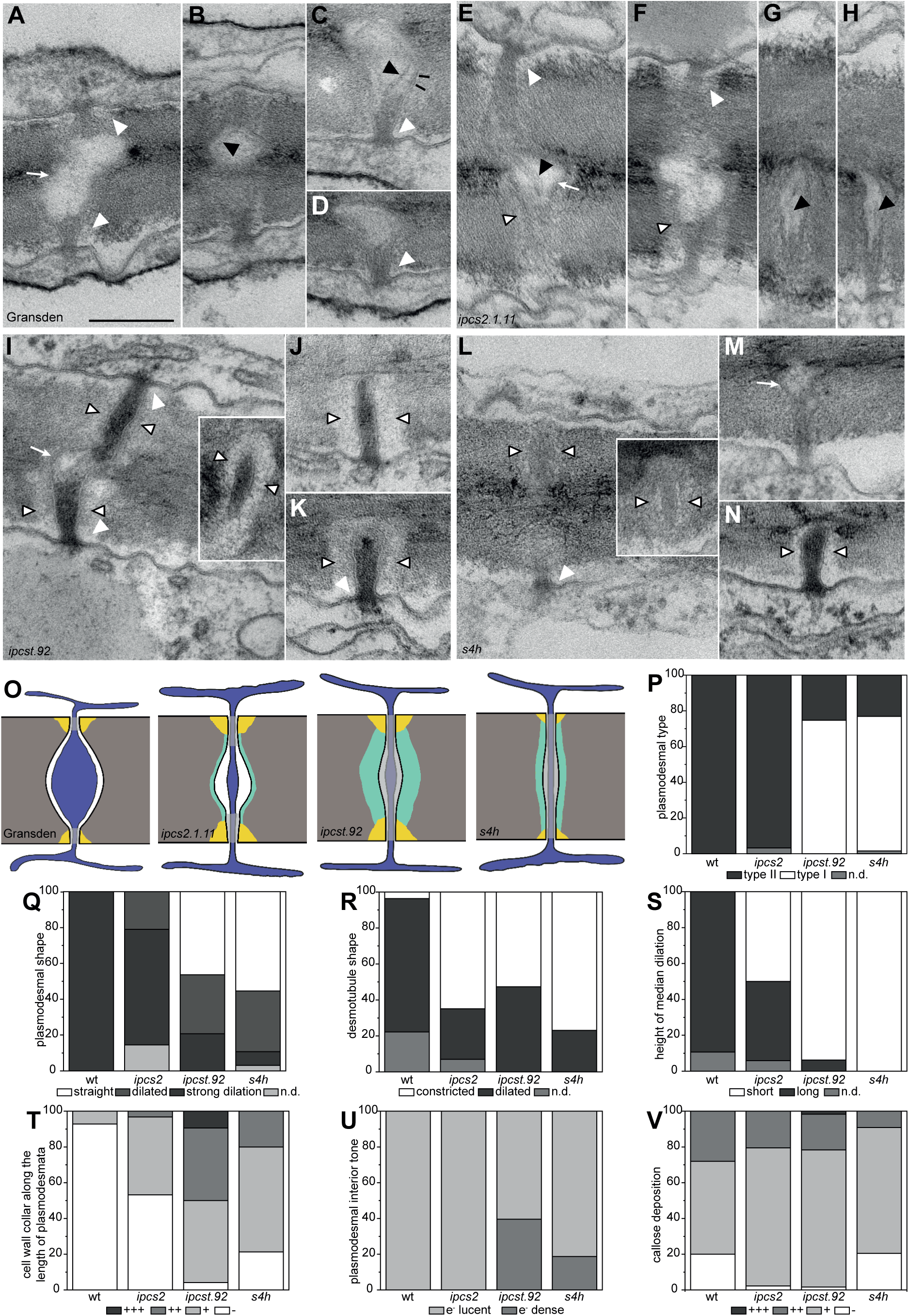
GIPC deficiencies are associated with changes to plasmodesmal ultrastructure. Transmission electron micrographs (**A-N**), schematic drawings (**O**), and quantification (**P-V**) showing the typical structures of plasmodesmata observed in anticlinal cell walls of mature phyllids. Micrographs show the wild-type Gransden (**A-D**), the *ipcs2.1.11* single mutant (**E-H**), the *ipcst.92* triple mutant (**I-K**), and the *s4h* mutant (**L-N**). The scale bar shown in **A** represents 200 nm in all TEM images. White arrowheads point at callose deposits constricting the neck regions; white arrows mark strong median dilations; black arrowheads point at the membranes of dilated (**B,C**) or constricted (**E,G,H**) desmotubules; open arrowheads indicate electron-lucent cell wall collars; and black lines in **C** point at rarely visible spoke-like proteins spanning the cytosolic sleeve. In the schematic drawings (**O**) all membranes are shown in black, the cell wall is depicted in grey, the ER is illustrated in dark blue, white shows the cytosolic sleeve between the plasma membrane and the central desmotubule. A grey overlay marks regions in which plasmodesmal substructures could not clearly be seen (i.e. necks and type I plasmodesmata). Yellow colour indicates callose deposits at the constricted plasmodesmal orifices, and aquamarine depicts electron-lucent cell wall collars surrounding the plasmodesmata. Graphs in (**P-V**) depict the percentages of plasmodesmata showing distinct traits of morphological characters. Only those plasmodesmata in which the indicated character was visible in the plane of section were taken into account, with **R** (desmotubule shape) referring only to the type II (-like) plasmodesmata in the respective line, and (**S**) (median dilation height) considering only those plasmodesmata showing strong median dilations.

All plasmodesmata observed in mature phyllids of the wild-type Gransden (Figure 4A-D, O, n=28) were classified as type II(-like) plasmodesmata (Figure 4P), since the desmotubule and the cytosolic sleeve could be clearly discerned in the median plasmodesmal regions, which always had a strongly dilated shape (Figure 4Q). In contrast to many characterized angiosperms, the desmotubule was also usually dilated in this region (Figure 4R), and spoke-like structures linking the desmotubule membrane to the plasma membrane could sometimes be observed (Figure 4C, indicated by black lines). The median dilations of the wild-type plasmodesmata were always long (Figure 4S) and covered more than half of the cell wall thickness. Cell wall collars surrounding the median region of the plasmodesmata were occasionally observed, in 7 % of the wild-type plasmodesmata (Figure 4T). The interior of the cytosolic sleeves and desmotubules were electron-lucent (Figure 4U), and short, constricted, electron-dense neck regions at the plasmodesmal orifices were surrounded by moderate callose deposits (Figure 4V). Thus, plasmodesmal characters of wild-type Gransden phyllids resemble those described previously for mature wild-type Reute phyllids (Wegner and Ehlers, 2024). However, at 0.158 plasmodesmata/µm^2^ cell wall (Table 1), the observed plasmodesmal density was substantially lower than in the Reute line (0.744 plasmodesmata/µm^2^ cell wall (Wegner and Ehlers, 2024)).

**Table 1:** Plasmodesmal densities (ρ(PD)) of the different GIPC-mutants based on plasmodesmal counts via TEM. The densities (PD/µm² cell wall) were calculated as described in (Wegner and Ehlers, 2024) based on the total number of plasmodesmata and the total cell wall length (µm). These values were gathered from at least 82 walls per genotype, which were observed in serial sections of one or two (*s4h*) biological replicates per genotype.

| | nWalls | nPD | PD/wall | total wall length | PD/$\mu\text{m}$ | $\rho(\text{PD})$ |
| --- | --- | --- | --- | --- | --- | --- |
| *Gransden* | 96 | 28 | 0.292 | 1,269.494 | 0.022 | 0.158 |
| *ipcs2.1.11* | 82 | 62 | 0.756 | 799.772 | 0.078 | 0.620 |
| *ipcst.92* | 90 | 93 | 1.033 | 915.258 | 0.102 | 0.813 |
| *s4h* | 166 | 64 | 0.386 | 1,699.678 | 0.038 | 0.301 |

In the *ipcs2.1.11* single mutant, plasmodesmal structure was moderately affected (Figure 4E-H, O, n=62). Observed plasmodesmata were still consistently type II (Figure 4P), with visible internal substructures and dilated median regions (Figure 4Q). However, in 21 % of the plasmodesmata, the median region was not as strongly dilated as in the wild type. *ipcs2.1.11* plasmodesmata also differed in that their desmotubules were mostly constricted (in 65 % of type II(-like) plasmodesmata) (Figure 4R). Additionally, *ipcs2.1.11* plasmodesmata were often surrounded by narrow, electron-lucent cell wall collars (in 47 % of plasmodesmata) (Figure 4T; wall collars depicted by open arrowheads in Figure 4E-N). While callose deposits at the neck regions were only slightly stronger than with the wild type (Figure 4V), the dilated median region did not cover half of the cell wall thickness, with 50 % of those plasmodesmata showing strong dilations (Figure 4S). The determined plasmodesmal density of 0.620 plasmodesmata/µm^2^ cell wall (Table 1) was about four times higher than in the wild-type Gransden from this study, and comparable to the published data for wild-type Reute.

An even higher density of 0.813 plasmodesmata/µm^2^ cell wall (Table 1) was found in the *ipcst.92* triple mutant (Figure 4I-K, O, n=87), whose plasmodesmata differed clearly from the wild-type Gransden, since 75 % (Figure 4P) failed to undergo the transition from type I into type II(-like) and did not show internal substructures. Among the few type II(-like) plasmodesmata, 53 % had a constricted desmotubule (Figure 4R). 46 % of the *ipcst.92* triple mutant plasmodesmata had a straight morphology with an almost constant diameter over their entire length, and did not develop any dilations in the median regions of the cell wall (Figure 4Q). Only 21 % of the *ipcst.92* plasmodesmata had strong median dilations resembling those of the wild type (Figure 4Q), but in most (94 %) of these, the median dilations did not cover half of the cell wall thickness (Figure 4S). Neck regions exhibited only slightly enhanced callose deposits (Figure 4V). One of the most striking features of *ipcst.92* triple mutant plasmodesmata was an electron-dense interior observed with 40 % of plasmodesmata (Figure 4I, K, U). This dense interior is often seen in young type I plasmodesmata (Nicolas et al., 2017). Additionally, electron-lucent cell wall collars were observed encasing 96 % of the analyzed plasmodesmata in *ipcst.92* (Figure 4I-K, T), a striking feature that was rare and far more subtle in the wild type (Figure 4T).

Plasmodesmata of the *s4h* mutant (Figure 4L-N, O, n=65) largely resembled those of *ipcst.92*, with 75 % keeping type I morphology in the mature state (Figure 4P). A constricted desmotubule was found in 77 % of the rare type II(-like) plasmodesmata (Figure 4R). The majority of plasmodesmata in the *s4h* mutant (55 %) had a straight morphology without any median dilation, and only 8 % of mutant plasmodesmata had strong median dilations (Figure 4Q), none of which covered half of the wall thickness (Figure 4S). Neck regions had slightly smaller callose deposits than the wild type (Figure 4V). An electron-dense interior was occasionally observed with 19 % of plasmodesmata (Figure 4U), and moderate electron-lucent cell wall collars were found around 79 % of the analyzed plasmodesmata (Figure 4T). An additional peculiarity of the *s4h* mutant was the low electron density of the plasma membrane; this is seen in plasmodesmata (Figure 4L, M), but also generally throughout the plasma membrane (Figure 3M, N). Further, in contrast to both *ipcs* mutants, the plasmodesmal density of the *s4h* mutant was only 0.301 plasmodesmata/µm^2^ cell wall (Table 1) and thus reaches only twice the value observed with the wild-type Gransden.

In summary, mutants with severe shifts in GIPC content failed to form normal type II(-like) plasmodesmata, and instead exhibit type I-like plasmodesmata with electron-dense interiors. Formation of dilated median plasmodesmal regions was impaired in all mutants, with desmotubule dilation being especially impaired relative to the wild type. The severity of the dilation phenotype increased with the overall strength of the mutant phenotype. Instead of median dilations, pronounced cell wall collars were visible around the mutant plasmodesmata.

### Incomplete cell division in *ipcst.92* allows free cytosolic and ER luminal movement between adjacent cells

To investigate the consequences of incomplete cell division and plasmodesmal structure in the mutants, we bombarded mature gametophores with fluorescent markers and observed and quantified intercellular movement of the fluorophores. Expression constructs with free eYFP (27 kDa) for cytosolic tracking and Sp-mCer-KDEL (27 kDa) for luminal ER were bombarded into all four genotypes co-cultivated on the same plates, and mature phyllids were used for imaging 2-3 days post-transformation (Figure 5A-D). In the wild type (Figure 5A, E), eYFP fluorescence was restricted 79 % of the time (15/19) to a single bombarded cell. The other 21 % of bombardment events led to some eYFP fluorescence in 1-3 additional cells adjacent to the bombarded cell. This could possibly be attributed to a relatively younger developmental stage of these cells and, consequently, a higher SEL of their plasmodesmata in these particular bombardment events (Wegner and Ehlers, 2024), or random, non-targeted leakage. In contrast, Sp-mCer-KDEL was always restricted to the bombarded cell (Figure 5A, F), fitting with previous observations that the desmotubule lumen does not allow for the transport of macromolecules larger than 3-10 kDa (Crawford and Zambryski, 2000; Li et al., 2021). For both *ipcs2.1.11* and *s4h*, as in wild type, the Sp-mCer-KDEL marker was not mobile between adjacent cells, except for one exceptional bombardment event in *ipcs2.1.11* (Figure 5F). With respect to cytosolic movement, wild type-like patterns were observed for *ipcs2.1.11* (Figure 5B, E), with 89 % (16/18) of bombardments producing only a single fluorescent cell, and the remaining events leading to weak fluorescence in 2-4 adjacent cells. In the tiny *s4h* phyllids, it was difficult to identify healthy bombarded cells with healthy neighbouring cells, which could be analyzed without interference from autofluorescence. In the 6 bombardments analyzed, eYFP did not diffuse to adjacent cells (Figure 5D, E). Due to the low replicate number and substantial morphological differences between *s4h* and all other genotypes analyzed, however, we cannot infer that there is any less cytosolic movement in this mutant compared to wild type. Indeed, it is counter-intuitive that there was not more cytosolic movement observed in the *s4h* mutant compared to the wild type; since the *s4h* plasmodesmata look like type I, a high SEL would have been expected that should regularly allow eYFP transport (Nicolas et al., 2017; Wegner and Ehlers, 2024).

**Figure 5:**
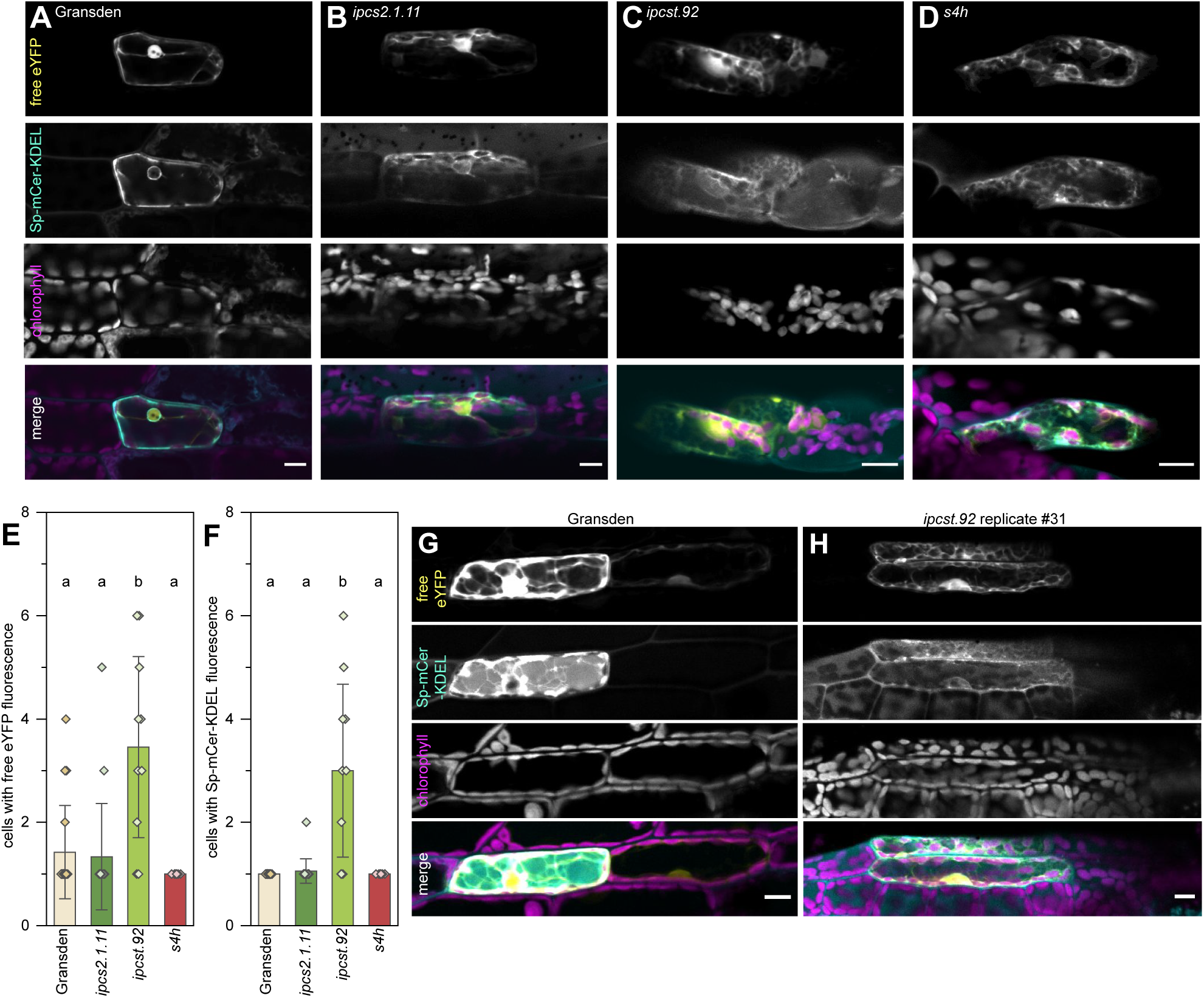
Intercellular connectivity is altered in *ipcst.92* mutant phyllids. (**A-D**) Representative images of free eYFP and Sp-mCerulean-KDEL expression and diffusion after bombardment in wild-type Gransden (**A**), *ipcs2.1.11* (**B**), *ipcst.92* (**C**), and *s4h* (**D**) Scale bars are 10 µm. (**E**,**F**) Quantification of cells exhibiting (**E**) free eYFP and (**F**) Sp-mCerulean-KDEL fluorescence per bombardment event. The bombarded cell is counted as n=1. Only bombardments in which both fluorophores were expressed were included. Bars are averages of 6-19 replicates with standard deviation, letters indicate significance at p < .05 determined by one-way ANOVA with Tukey’s *post-hoc* test. (**G**,**H**) Representative images of low eYFP diffusion in wild type, presumably through plasmodesmata (**G**), and strong eYFP and mCerulean diffusion in *ipcst.92* replicate line #31, presumably due to transport through incomplete cell walls (**H**). The replicate triple mutant line #31 was used to confirm intercellular motility was not unique to the *ipcst.92* mutant allele. Scale bars are 20 µm.

In *ipcst.92*, of 11 bombardment events, only 18 % (2/11) produced just a single eYFP fluorescent cell, while for the remaining bombardments there was diffusion to 1-5 adjacent cells (Figure 5C, E, H). Further, unlike all other genotypes tested, in *ipcst.92* the cells adjacent to the bombarded cell almost always exhibit fluorescence of not only cytosolic eYFP, but also the Sp-mCer-KDEL marker (Figure 5C, F, H). The appearance of eYFP-fluorescent cell clusters in *ipcst.92* was also distinct from the other genotypes in that the clusters were generally uniformly bright with eYFP, to the extent that the bombarded cell could not be distinguished from its neighbours (Figure 5H). In the other genotypes, weak eYFP fluorescence in cells adjacent to the bombarded individual cell could only be visualized by either adjusting microscope settings or enhancing image brightness to the point that fluorescence signal from the bombarded cell was saturated (Figure 5G). We infer that in wild type, eYFP is able to pass through plasmodesmata of adjacent cells, but with a limited capacity that is dependent on the SEL of plasmodesmata, the developmental stage of the bombarded gametophore colony, phyllid, and individual cell. In the *ipcst.92* mutant, in contrast, there was unregulated cytosolic eYFP movement within fluorescent cell clusters. Together with the intercellular spread of the mCerulean fluorophore, which is well above the size of other desmotubule-mobile molecules, these results suggest that there is free, non-plasmodesmal connectivity of both cytosol and ER lumen in *ipcst.92*, most likely via the incomplete anticlinal cell divisions observed in Figure 2L, N.

### Modified plasmodesmal structure in GIPC-deficient mutants does not abolish their general capacity for intercellular cytosolic transport

We observed no transport of eYFP and Sp-mCer-KDEL molecules with the *s4h* mutant, so the question arose, whether these plasmodesmata are functional at all. This question was also provoked by the strong structural abnormalities of all mutant plasmodesmata observed by TEM. To test whether smaller molecules can move cell to cell, we performed FRAP experiments on phyllids stained with the small fluorescent dye carboxyfluorescein diacetate (CFDA, broken down to and sequestered as carboxyfluorescein, 376 Da, upon cellular uptake). We compared CF FRAP in parenchyma cells from phyllids dissected from the gametophore tips. In all genotypes, we could find both cells that did not recover fluorescence, even 10 or 20 minutes after photobleaching, and cells that partially or fully recovered fluorescence more or less quickly within this observation period (Figure 6, A-D).

**Figure 6:**
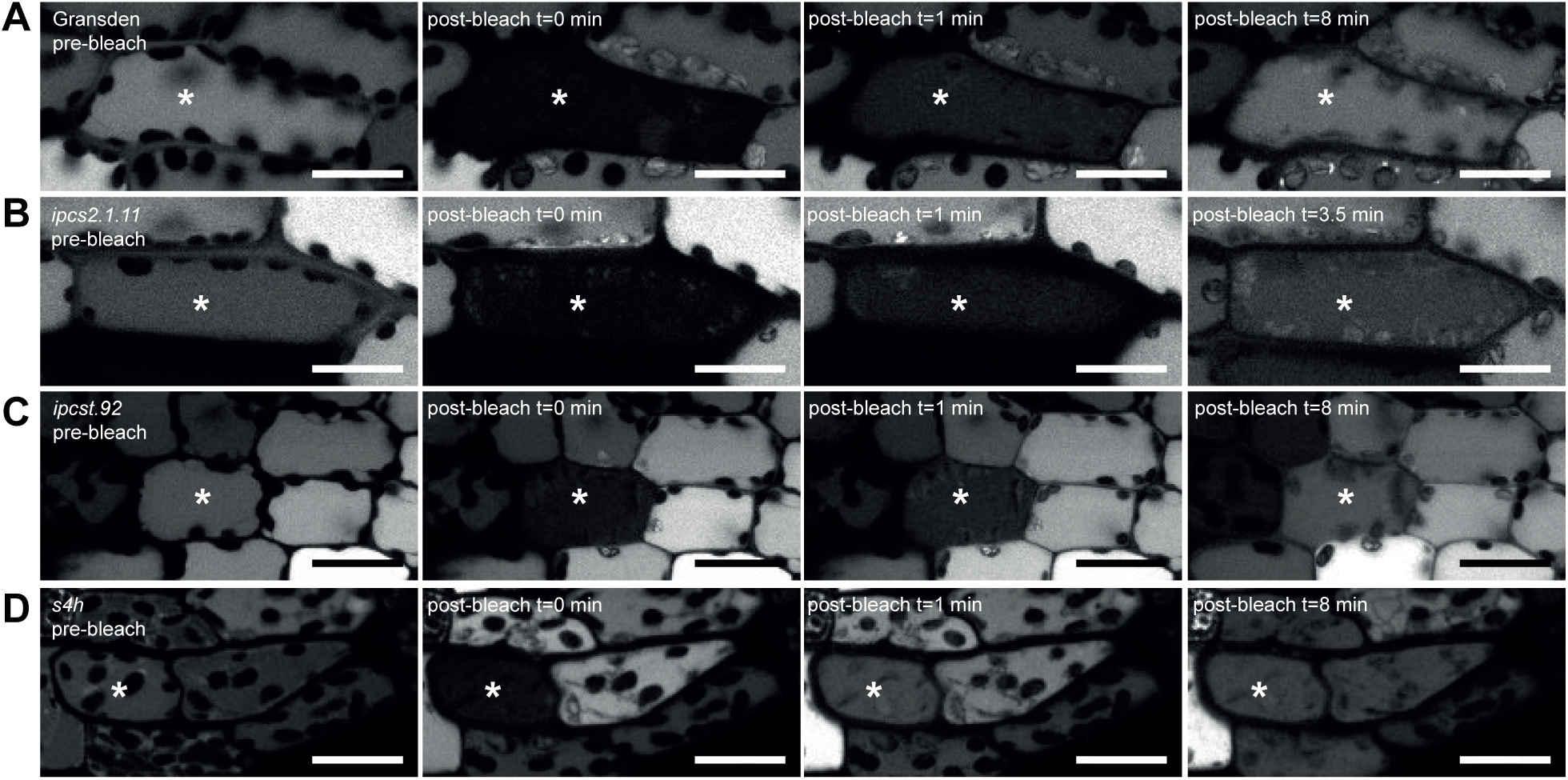
FRAP of cytosolic carboxyfluorescein in phyllids of wild type and GIPC-deficient mutants over time (t). **A** wild type B *ipcs2.1.11* C *ipcst.92* D *s4h*. Asterisks mark the bleached cells. All scale bars 20 µm.

The observation that CF dye movement was similar to wild type is especially meaningful for *s4h*, as we had not observed eYFP fluorophore movement in this genotype after bombardment. Redistribution of CF after bleaching demonstrates that despite strong plasmodesmal structural defects and no observation of macromolecular transport, the plasmodesmata retain some functionality.

For the *ipcst.92* mutant, the CF dye redistribution after photobleaching could be tentatively associated with plasmodesma-mediated transport, in contrast to the eYFP spread after biolistic bombardment. We identified cells in *ipcst.92* which recovered fluorescence (Figure 6C), but which were less likely to be cytoplasmically connected to adjacent cells due to incomplete cell division by pre-examining their cell walls in Z-stacks of the dye-stained phyllids (Supplementary Figure S3A). With this approach the incomplete cell walls, and intercellular motility of cytosol and organelles were even more obvious (Supplementary Figure S3A, Supplementary Video SV1). As observed previously with light and electron microscopy of thin-sectioned phyllids (Figure 2L, N; Figure 3A, B), here we could confirm that cell wall gaps and attachments sites showed no consistent orientation or placement relative to the whole phyllid. Cells with incomplete cell walls had a characteristic response to photobleaching: local loss of fluorescence was never complete, and rapidly recovered within a minute after bleaching (Supplementary Figure S3B). A quantitative comparison of FRAP recovery rates was not realistic due to developmental differences among the mutants; however, the fact that recovery could simply occur in selected cells of both *ipcst.92* and *s4h* indicates that their plasmodesmata are generally functional and allow the transport of small molecules, despite structural deviations from the wild type.

## Discussion

### Cell division and membrane morphology are impacted when GIPC synthesis is manipulated in *P. patens*

We isolated *P. patens* mutants with GIPC deficiencies including weak and strong overall reductions, and dramatic shifts in composition. One of the most severe deficiencies we observed in overall GIPC content in *P. patens* was in the *ipcs* triple mutant allele *ipcst.92*, which shows especially strong reductions in Hex- and Hex-Hex-GIPCs, but no significant increase in free ceramide levels. We previously reported gross morphological defects in this mutant, and here show that this dwarfism is coincident with a drastic reduction in cell size and especially cell number per phyllid. We infer a reduced frequency of cell division. Further analysis with light and electron microscopy linked this phenotype to incomplete anticlinal cell divisions. Cellular bombardment and FRAP assays demonstrated that the openings within anticlinal cell walls allow for free movement of cytosol, ER, and other organelles between adjacent cells. In light of this dramatic ultrastructural defect, it is remarkable that the *ipcst.92* mutant is not only viable, but also produces differentiated cells: it can establish protonema and gametophores, including phyllids with hadroms, albeit with weaker structural definition than in the wild type. Notably, differentiation from gametophores to protonema in *ipcst.92* is slow; beginning with homogenization of gametophores, the production of homogenous protonema requires 4-6 months of weekly cultivation.

Observation of defects in cell division associated with sphingolipid deficiency is not without precedent. Previous work described Fumonisin B1 (FB1)-treated *A. thaliana* roots, which are impaired in the synthesis of VLCFA-containing ceramides that normally channel into the assembly of GIPCs (Molino et al., 2014). FB1-treated plants exhibit complex shifts in their sphingolipid profiles, however two of the more substantial shifts are: (1) an overall increase in free ceramide levels, and (2) a flattening of the chain length profile of the fatty acid moieties of sphingolipids, namely an increase of ceramides with 16 carbon LCFAs that could partially compensate for diminished VLCFA-containing ceramides (Markham et al., 2011). FB1 treatment was linked to a mild reduction in cell size, and a more substantial ∼25% reduction in cell number, within the root division zone in *A. thaliana* (Molino et al., 2014). Although *P. patens ipcs* mutants do not show substantial shifts in the fatty acid moiety chain length profile of GIPCs specifically, there is likely an overall reduction in VLCFA content for membrane lipids as a whole due to the reduction in GIPC content. Therefore, we cannot distinguish whether the cytokinesis defects in *ipcst.92* are due to reduction in GIPCs, or reduction in VLCFAs. Further, it is unknown whether the cell division phenotype of *ipcst.92* is directly caused by sphingolipids, or if it could be mediated by downstream effects on the overall lipid content of cellular membranes. A mechanism whereby GIPCs (as VLCFA-containing sphingolipids) could affect cytokinesis was previously proposed (Molino et al., 2014), namely that VLCFAs stabilize highly-curved membranes that form upon vesicle fusion, which is of particular importance during cell plate formation. In both our *P. patens ipcst.92* mutant and in FB1-treated *A. thaliana* roots, plate formation appears to be aborted during late cytokinesis, after substantial vesicle fusion must have already occurred. If our model is correct, this implies there is sufficient VLCFA and/or GIPC content in these membranes to support the initial stages of vesicle fusion, but that at some point it can no longer support stable expansion of the cell plate. Overall, this hypothesis and model parallels observations in *S. cerevisiae* that VLCFA-containing lipids are essential for formation of the highly-curved membranes of nuclear pores (Schneiter et al., 2004).

Similar phenotypes to *ipcst.92* have were also observed in *A. thaliana pi4kβ1 pi4kβ2* double mutants deficient in PI4P synthesis, that is, aborted cytokinesis producing multinucleate cells (Lin et al., 2019). Additionally, a different, but related phenotype was reported by (Lebecq et al., 2022; Lebecq et al., 2023) who showed that localized depletion of phosphatidylinositol 4,5-bisphosphate [PI(4,5)P_2_] by the phosphatase SUPPRESSOR OF ACTIN9 (SAC9) is required at the leading edge of the phragmoplast; in *sac9* mutants, ectopic accumulation of PI(4,5)P_2_ leads to cell plate branching. The similarity of *P. patens ipcst.92* to *A. thaliana pi4kβ1 pi4kβ2* specifically is perhaps surprising in light of recent analysis of *Atipcs1/2* amiRNA lines, which revealed PIP phospholipase activity is dependent on the presence of VLCFA-containing sphingolipids in *A. thaliana* (Ito et al., 2021). This study demonstrated that reduced levels of VLCFA-containing sphingolipids produced increased PI4P at the TGN, suggesting an inverse relationship between GIPC and PI4P accumulation. The similarity of our GIPC-deficient phenotypes to PI4P-deficient phenotypes is in apparent opposition with the notion that GIPC accumulation is antagonistic to PI4P accumulation. However, we were unable to measure phosphoinositide levels in *P. patens* for the present study, and our approach for GIPC detection is insensitive to subcellular localization. More precise analysis of the spatiotemporal changes in sphingolipid and phosphoinositide content in our mutants may present a more harmonious model of how these lipid classes interact. Additionally, it is not inconceivable that there may be differences between *A. thaliana* and *P. patens* in this respect, considering substantial differences between cytokinesis in these two models, reviewed in (Buschmann and Zachgo, 2016; Buschmann and Müller, 2019), and the fact that the vascular and bryophyte lineages diverged over 450 million years ago.

The *s4h* mutant has a complex and unique metabolic phenotype: *s4h* accumulates sphingolipids with 18:0;2 in place of 18:0;3, and has strong accumulations of B-series Hex-Hex-GIPCs and Hex-HexNAc-GIPCs. Among the visual phenotypes observed in *s4h*, we were especially interested in the loss of orientation of cell division within phyllids. This observation together with the overall morphological abnormalities of the *s4h* mutant suggest that it may have partially lost the capability to define apical-basal polarity. Another interesting aspect of the *s4h* phenotype is the tufted aggregates observed here and in previous studies (Gömann et al., 2021; Steinberger et al., 2021). The aggregates observed with various imaging techniques must represent multiple different chemical accumulations. The aggregates likely include callose, as suggested by aniline blue-stained accumulations adjacent to cell walls seen with epifluorescence microscopy, and electron-lucent layers within the depositions attached to cell walls in TEM images. Callose hyperaccumulation and mis-localization is reminiscent of persistent callose at the maturing cell plate of *A. thaliana* observed by Molino et al. (2014) upon FB1 treatment, though the phenotype we observed is more severe. The red-coloured material in tufted aggregates viewed by light microscopy, and the electron-dense material in tufted aggregates observed by TEM, is not callose. This would be in conflict with their pigmentation and osmium tetroxide staining, and with the spatial separation of these features. The chemical composition of these structures remains ambiguous. Fragmentation of chloroplasts was frequently observed in *s4h* with basic light microscopy, as well as patchy CF fluorescence suggesting inconsistent hydrolysis of non-fluorescent CFDA to produce fluorescent CF. Both observations suggest cell death in *s4h* phyllids, with a patchy, cell-autonomous distribution. Whether and how cell death may be correlated with development of tufted aggregates was not investigated in the present study, and remains an essential question for understanding this mutant phenotype.

The plasma membrane detachment phenotypes we observed in all mutants could support the notion that GIPCs impact the physical connection of the plasma membrane and cell wall. These phenotypes could also, alternatively or additionally, be driven by accumulation of excess plasma membrane material in the mutants, such that the plasma membrane area exceeds the inner cell wall area. Vesicular trafficking is likely impacted by GIPC deficiencies, given GIPC enrichment along the secretory pathway and the fact that GIPCs impact membrane physical properties: they increase membrane thickness due to their incorporation of VLCFAs (Levine et al., 2000), and increase lipid ordering due to their incorporation of saturated, hydroxylated LCB and VLCFA moieties that hydrogen bond with each other and free sterols in the membrane (Grosjean et al., 2015). Further, as discussed above, VLCFA-containing GIPCs are expected to be required for vesicle fusion during cytokinesis. It seems reasonable to expect that other dynamic processes involving plasma membrane budding and fusion would be impacted by sphingolipid content, including secretion and endocytosis, and this in turn would affect the staining properties and physical appearance of the plasma membrane.

### Plasmodesmal development is impacted by GIPC deficiencies in *P. patens*

The *s4h* and *ipcst.92* mutant plasmodesmata did not undergo a full type I-to-type II(-like) transition (Nicolas et al., 2017; Wegner and Ehlers, 2024). Internal plasmodesmal substructures did not become visible along the length of these severely affected plasmodesmata, there was no clear spatial separation of the desmotubule and the plasma membrane by a visible cytosolic sleeve, they regularly exhibited an electron-dense interior, and the widening of the plasmodesmal diameter and the formation of significant median dilations was lacking in most plasmodesmata of these mutants. Striking cell wall collars were observed instead. It could be speculated that the developmental defects of the severely affected mutants’ plasmodesmata would impair their quantitative transport capacities, that is, the volume flow of even small molecules. Enlarged cytosolic sleeves and/or enlarged median dilations characteristic of type II(-like) plasmodesmata would otherwise contribute to plasmodesmal widening. This is thought to sustain transport rates when plasmodesmal channels become elongated during thickening growth of the wall: Modelling predictions showed that larger cytosolic sleeves and median dilations would be favoured in thicker walls to increase plasmodesmal permeation efficiency (Deinum et al., 2019). Lack of these plasmodesmal modifications and reduced symplastic transport rates might contribute to the dwarf phenotypes of all three mutants. Notably, in *A. thaliana*, multiple C2-domains and transmembrane region proteins (MCTPs) have been shown to act as plasmodesmata-specific ER-plasma membrane tethers interact with PI4P, contribute to plasmodesmal formation and function (Li et al., 2023; Pérez-Sancho et al., 2023), and MCTP-deficient mutants show dwarfism and aberrant cell division phenotypes (Brault et al., 2019) that are similar to *s4h*. MCTPs have been identified in the *P. patens* plasmodesmal proteome and confirmed to localize to plasmodesmata via confocal microscopy (Gombos et al., 2023). It will certainly be of interest to functionally characterize MCTPs in *P. patens* and to test whether their localization, and the localization of other plasmodesmal and plasma membrane proteins, are disrupted in *s4h*.

In addition to the plasmodesmal structural defects in our mutants, we observed two-fold increased plasmodesmal density in *s4h*, and four- to five-fold increased plasmodesmal densities in *ipcs2.1.11* and *ipcst.92* compared to the wild type. One explanation that could account for both the observed plasmodesmal structural defects and their increased density in our mutants is that both of these phenotypes reflect that the mutant phyllids are altogether not fully differentiated, perhaps displaying some default state, or incomplete gametophore-specific programming. As discussed further below, *P. patens* protonemal cell walls are characterized by a very high density of type I plasmodesmata, and lack type II(-like) plasmodesmata (Gombos et al., 2023). Similarity of wild-type protonema to the mutant phenotypes observed in gametophores supports the notion that the mutants may simply be less differentiated by normal gametophore developmental programming. Moreover, type II(-like) plasmodesmata can be expected to mediate a highly controlled, targeted exchange of macromolecular signals whereas type I PD would allow their unhindered non-specific transport by diffusion (Crawford and Zambryski, 2001; Ehlers and Westerloh, 2013; Nicolas et al., 2017). Thus, lack of type II(-like) PD formation in the *s4h* and *ipcst.92* mutants could explain their severe developmental phenotypes, in particular their impaired ability to undergo the developmental switch from simple protonema- to complex gametophore growth and their problems in tissue differentiation. Clearly, in *ipcst.92*, these issues would be compounded by incomplete cell division.

A second model to explain the changes in plasmodesmal density and structure is that the cytokinetic defects, of *ipcst.92* and *s4h* in particular, cause the plasmodesmal phenotypes: As it has been demonstrated that primary plasmodesmal deposition is determined by ER strands becoming trapped and integrated in the expanding cell plate (Hepler, 1982; Li et al., 2023), perhaps the cytokinetic defects are somehow responsible for the higher frequency of ER “trapping” in the plate, and/or the stabilization and retention of desmotubules. Overall, it seems most likely to us that the plasmodesmal phenotypes observed in these sphingolipid-deficient mutants are at least partially caused by broader developmental and cytokinetic defects.

We could also make inferences regarding the SEL of mutant plasmodesmata from the FRAP and biolistic assays. Due to the ambiguity of these results in the *ipcst.92* mutant introduced by its incomplete cell divisions, we will limit our discussion to the *ipcs2.1.11* and *s4h* mutants. Similarly to the Gransden wild type, both the *ipcs2.1.11* with weakly dilated type II-like plasmodesmata, and even *s4h* mutant with narrow and straight type I plasmodesmata mediated the fast exchange of CF(DA) (376 Da) between mature phyllid cells within minutes after bleaching. This suggests that the shape of plasmodesmata and their desmotubules, which are dilated in wild-type plasmodesmata, less dilated in *ipcs2.1.11*, and mostly constricted in *s4h*, did not have a substantial impact on the general capacity for transport of small molecules. For *ipcs2.1.11,* slow intercellular exchange of eYFP (27 kDa) was also occasionally observed, similar to the Gransden wild type. A comparison can be made to *P. patens* protonema, which exclusively form straight plasmodesmata at very high density (Gombos et al., 2023). Movement of Dendra2 (26.1 kDa) between protonema cells, presumably *via* type I plasmodesmata, could be observed within 15 min, whereas between mature *P. patens* phyllid cells, which are interconnected by type II-like PD (Wegner and Ehlers, 2024), movement was only observed after 9.5 h over 1-2 cells (Kitagawa and Fujita, 2015). The slow spread of Dendra2 into neighboring phyllid cells indicates that while macromolecular transport rates are drastically reduced in this tissue compared to protonema cells and young phyllids with type I plasmodesmata (Wegner and Ehlers 2024), transport of larger molecules is not entirely prevented by a strict size exclusion limit in mature phyllids (Kitagawa and Fujita, 2015). At this point, it is unclear whether this slow macromolecule transport over hours and days has a physiological impact. Regardless, we infer that developmental control of the plasmodesmal SEL is exerted in regions of the plasmodesmata largely unaffected by the transition from type I to type II and by these mutations, and instead presumably occurs *via* molecular changes in the neck regions at the plasmodesmal orifices. The persistent finding of occasional, very slow macromolecule (Dendra2 and eYFP) transport through plasmodesmata raises the question whether the SEL is indeed an exclusively qualitative and definite parameter; it could be that this reflects random leakage that is unavoidable during targeted macromolecule transport.

### *P. patens* mutants are complementary tools for deciphering complex developmental phenotypes

Plant development is intricate and is easily derailed past the point where processes can be cleanly analyzed and deciphered when using blunt mutagenesis approaches. Plant species that have relatively simple organ structure and developmental programming are therefore uniquely valuable, and the moss *P. patens* is in this sense ideal: throughout its predominantly haploid life cycle, almost all organ structures are only a single cell layer thick, including the filamentous protonema, rhizoids, and gametophore phyllids (Cove et al., 2006; Vidali and Bezanilla, 2012). Here, we have described functions for GIPCs in *P. patens* growth and development, with severe and specific deficiencies in cytokinesis linked with specific manipulations of GIPC synthesis. These findings are complementary to previous work in *A. thaliana*, and support the growing body of evidence that VLCFA-containing GIPCs are required for normal endomembrane dynamics, likely specifically due to a role in supporting vesicle fusion as previously proposed (Molino et al., 2014). These congruent results in *P. patens* additionally suggest that GIPC functions could be conserved across embryophytes. We also observed that GIPCs impact plasmodesmal development, likely at least in part as an effect of cytokinetic and gross developmental defects. These mutants and the phenotypic, biolistic, and FRAP assays developed here can be used to further explore the precise roles of sphingolipids in plant development.

## Methods

### Cultivation

*P. patens* ecotype Gransden 2004 was obtained from the International Moss Stock Center (IMSC; https://www.moss-stock-center.org/en/), strain 40001. Plants were maintained under long-day conditions (16 h light/8 h dark, at 25 °C /18 °C) with 105-120 µmol m^−2^ s^−1^ radiation. Filamentous protonema tissue was cultivated on BCD-AT medium (1 mM MgSO_4_, 1.84 mM KH_2_PO_4_, 10 mM KNO_3_, 45 µM FeSO_4_, 5 mM ammonium tartrate, 1 mM CaCl_2_, Hoagland’s trace elements, 0.55 % plant agar) overlaid with sterile cellophane discs, and the cultures were propagated by disruption in sterile tap water with an IKA ULTRA-TURRAX homogenizer every 1-2 weeks. Development of gametophores was induced by transferring small pieces of tissue to BCD medium (BCD-AT lacking ammonium tartrate). Gametophores were sub-cultivated every 4-6 weeks (Maronova and Kalyna, 2016). The *s4h* mutant was generated *via* homologous recombination as described (Gömann et al., 2021), with the entire coding sequence replaced by a kanamycin selection marker. Generation of the *ipcs* mutant series with CRISPR/Cas9 was previously reported (Haslam et al., 2024); *ipcs2.1.11.* contains a single nucleotide insertion 76 bp from the ATG start of *IPCS2*, resulting in an early frame shift in the coding sequence. The *ipcst.92* mutant has frame-shift lesions in *ipcs1* (16 bp deleted 32 bp downstream from the ATG start) and *ipcs3* (16 bp deletion and 2 bp substitution, 189 bp downstream from the ATG start), while the *ipcs2* lesion is an out-of-frame 12 bp deletion, resulting in the substitution of IELTV>M, a putative alternative translational start. Our previous work demonstrated via allele replacement in *P. patens* and heterologous expression in *Saccharomyces cerevisiae* that this specific *ipcs2* lesion does impair IPCS2 activity.

### General growth phenotype analyses

Gametophore colony area and circularity were measured from two-month old plants. After ventilating plates under a sterile bench to reduce humidity and condensation, mutant and wild-type gametophores were photographed directly on covered petri dishes, using a Retiga R6 camera on an Olympus SZX12 stereomicroscope. Image analysis was carried out with ImageJ 1.53f5.1 (Schneider et al., 2012). Area was measured by selection of gametophores with the Wand tool and adjustment of selection tolerance. Gametophore circularity was calculated as C = 4 πA/P^2^ (C=circularity; A=area; P=perimeter) (Vidali et al., 2007; Vidali and Bezanilla, 2012).

Cell number per phyllid and cell size were measured from phyllids excised from mature gametophore colonies. Phyllids were imaged on an Olympus BX51 light microscope with a 10X objective, and a Retiga R6 camera. We tried to select phyllids #6-12 from the apex of each gametophore shoot for measurements. For counting total cell #/phyllid, we counted cells on one side of the hadrom and multiplied this number by two. Measurements were carried out manually with ImageJ 1.53f5.1.

Semi-thin sections of 0.5-1 µm thickness were cut from the median plane of fixed and resin-embedded phyllids with glass knives on a Reichert Om U2 ultramicrotome (Leica Microsystems GmbH, Wetzlar, Germany) for histological investigations. Sections were stained with 0.5 % crystal violet and observed under a Leica DM 5500 microscope B equipped with a Leica DFC 450 camera driven by the Leica Application Suite software (Version 4.3.0).

### Aniline blue staining and combined light and fluorescent microscopy

Live cell staining with aniline blue paired with light and epi-fluorescent microscopy was used to compare the localization and test for colocalization of different pigmented, stained, or fluorescent compounds observed in various studies of the *s4h* mutant. Live cell staining followed the protocol described by (Sankoh et al., 2024). Gametophore phyllids were then imaged on two separate Olympus BX51 light microscopes, one fitted with a Retiga R6 color camera, and another with a black and white Hamamatsu ORCA-Flash4.0 LT+ camera. An F46-001 ET-Set CFP filter cube was used for aniline blue and F41-007 HQ-Set Cy3 for autofluorescence.

### Biolistic transformation

Marker plasmids were bombarded into 4-6 week old gametophore colonies. Within experiments, different genotypes were cultivated on the same 5 cm Petri plate, and were therefore transformed with the same plasmid preparations, precipitated on the same microcarriers, and bombarded in the same shot. Free enhanced yellow fluorescence protein (eYFP, 27 kDa) was expressed from a pUC18-derived entry vector, driven by the 35S promoter. The ER marker plasmid was received from Prof. Ralf Reski, University of Freiburg; this consisted of the *P. patens* aspartic protease signal peptide fused to mCerulean (27 kDa) with C-terminal KDEL retention signal (SpCerKDEL), driven by the *P. patens ACTIN5* promoter, in a pJET1.2-based vector (Mueller and Reski, 2015).

Microcarriers were prepared and precipitated on macrocarriers as described (Müller et al., 2017), with minor modifications. Briefly, 60 mg of 1.0 µm gold particles (Biorad, #1652263) were suspended in 1 mL 70 % ethanol, washed in ultra-pure water, and finally suspended in 1 mL 50 % glycerol, for a final concentration of 60 mg/mL.

To prepare microcarriers sufficient for two bombardments, 4 µg of each plasmid to be transformed or co-transformed was used to coat a 10 µL aliquot of the 60 mg/mL gold suspension. The 10 µL aliquot of gold was first vortexed, the plasmid DNA added, and then sterile water, to bring the mixture volume up to 55 µL. Then 50 µL 2.5 M CaCl_2_ and 20 µL 0.1 M spermidine were added. The mixture was vortexed for 2 min then briefly spun down at low speed. The carriers were washed twice in 96 % ethanol, and finally re-suspended in 60 µL 96 % ethanol, then 2 X 30 µL were dispersed on macrocarriers (Biorad, #1652335) for bombardment. Gametophores were bombarded in a Bio-Rad PDS-1000/He Biolistic Particle Delivery System, with 900 psi rupture discs, and the vacuum pressure raised to 24 inHg before bombardment. After bombardment, gametophores were cultivated under normal growth conditions for two to three days before imaging.

### Symplastic motility assays

Images of bombarded cells of young, apical phyllids were captured on a Zeiss Axio Imager.Z2 LSM980 confocal laser scanning microscope, with a Plan-Apochromat 20X/0.8 M27 objective. eYFP was excited with a 514 nm diode laser at 0.8 % intensity with a 26 µm/0.9 AU pinhole, with emission detected between 525-569 nm. Chlorophyll and mCerulean were both excited with a 405 nm diode laser at 2.0 % intensity with a 26 µm/1.0 AU pinhole; chlorophyll fluorescence was detected between 645-713 nm, and mCerulean between 460-525 nm. Images were processed with ZEN3.2 and ImageJ 1.53f5.1.

### Carboxyfluorescein fluorescence redistribution (/recovery) after photobleaching (FRAP)

Whole *P. patens* gametophores were stained in aqueous 1 mM 5-(and-6)-carboxyfluorescein diacetate (CFDA, fluorescent and membrane impermeant upon hydrolysis to carboxyfluorescein, CF, 376 Da) for 60-90 min at RT, and subsequently washed four times in water. The gametophores were incubated in tap water overnight in the dark at room temperature to improve stain distribution. During this long incubation period, some CF was sequestered from the cytoplasm into the vacuoles.

FRAP was carried out with a Zeiss Axio Imager.Z2 LSM980 confocal laser scanning microscope, with a Plan-Apochromat 40X/1.4 oil immersion objective. CF was bleached 50 X at 2.52 s/frame with 100 % intensity of 405 and 488 nm diode lasers. The pinhole was 28 µm/0.88 AU. CF fluorescence was detected with the 488 nm laser at 1-3 % intensity, between 507-537 nm with a multi-alkali photomultiplier tube (PMT). Images were analyzed with ZEN3.2 and processed with Image J 1.53f5.1.

### Transmission electron microscopy

Sample preparation for light and electron microscopy followed the protocol of (Wegner and Ehlers, 2024). Whole gametophores were embedded in 2 % (w/v) low gelling agarose (type VII Sigma-Aldrich, Steinheim, Germany) and solidified agarose blocks were fixed in 50 mM phosphate buffer containing 2.5 % (v/v) glutaraldehyde (GA) at pH 7.23 for 2 h at RT and for another 2 h on ice. After rinsing in 100 mM buffer, postfixation was performed in 0.9 % (w/v) OsO_4_ in 0.1 M buffer at 4 °C overnight. Rinsing in water was followed by staining for 2 h with 0.5 % (w/v) aqueous uranyl acetate on ice, and dehydration in a graded ethanol series and propylene oxide. Samples were embedded in Spurr’s epoxy resin (adapted mixture; (Spurr, 1969)) and polymerized at 68 °C for 20 h.

Ultrathin sections (∼80 nm) were cut with a diamond knife, transferred onto formvar-coated single-slot copper grids, and contrasted with 2% uranyl acetate and lead citrate (Reynolds, 1963) for 12 min each. Sections were analyzed with an EM912AB TEM (Zeiss, Oberkochen, Germany) at 120 kV accelerating voltage under zero-loss energy filtering conditions. Micrographs were recorded with a 2k×2k dual-speed slow-scan CCD camera (SharpEye, TRS, Moorenweis, Germany) using the iTEM software package (OSIS). Figure plates were mounted with Corel PHOTO-PAINT (2021, Version 23.1.0.389, Corel, Ottawa, Canada) and in Adobe Illustrator CS6 Version 16.0.3, measurements were performed with ImageJ 1.53e (Schneider et al., 2012) and PD densities were determined according to (Wegner and Ehlers, 2024).

### Lipidomic analysis

Gametophores were harvested and lyophilized after one or two months of growth, dependent on the growth rate of the genotype.

#### Microsome enrichments

Microsome enrichments followed a published protocol (Abas and Luschnig, 2010). 10 mg lyophilized tissue was taken from each sample, and ground with stainless steel beads in a laboratory mixer mill MM400 (Retsch GmbH, Haan, Germany). The pulverized tissues were then suspended in a fractionation buffer consisting of 150 mM Tris-HCl (pH 7.5), 37.5 % sucrose, 7.5 % glycerol, 15 mM EDTA, 15 mM EGTA, and 7.5 mM KCl. The suspensions were vigorously mixed, then centrifuged at 1,500 g for 3 min at 4 °C. The supernatant was retained, and the pellet re-extracted twice with the fractionation buffer diluted first 0.75 X, then 0.67 X. All of the supernatants were pooled, diluted 1:1 with water, then spun at approx. 21,000 g for 2 hr. The pellets were washed, re-spun for 45 min, then finally sealed under argon gas, snap frozen in liquid nitrogen, and stored at −80 °C.

#### Lipid extractions

Total lipid extraction followed a published protocol (Herrfurth et al., 2021). Microsome pellets were re-suspended in 6 mL extraction buffer consisting of isopropanol:hexane:water in a 60:26:14 ratio, v:v:v. They were vortexed, sonicated, and shaken at 60 °C for 30 min. Cell debris was then spun down at 800 g for 20 min, at room temperature. The supernatant was dried under a nitrogen stream and finally resuspended in 800 µL tetrahydrofuran:methanol:water in a 4:4:1 ratio, v:v:v (TMW). The samples were stored at −20 °C, or processed directly.

#### Methylamine treatment

Methylamine treatment followed the protocols of (Markham and Jaworski, 2007; Herrfurth et al., 2021). Lipid extracts to be used for sphingolipid measurement were treated with methylamine to hydrolyze glycerolipids. This fraction was dried under a nitrogen stream, then re-suspended in 700 µL 33 % methylamine in ethanol (v/v) and 300 µL water. The samples were incubated at 50 °C for 1 h, and then dried again under nitrogen. The residue was finally re-suspended in TMW.

#### Ultra-high pressure liquid chromatography coupled with nanoelectrospray and tandem mass spectrometry (UPLC-nanoESI-MS/MS)

Sample injection of 2 µL was performed by an autosampler set at 18 °C and sample separation was conducted at a flow rate of 0.1 mL/min. The solvent system was composed of methanol:20 mM ammonium acetate (3:7, v/v) with 0.1 % acetic acid (v/v) (solvent A) and tetrahydrofuran:methanol:20 mM ammonium acetate (6:3:1, v/v/v) with 0.1 % acetic acid (v/v) (solvent B). A linear gradient was applied: start from 65 % B for 2 min; increase to 100 % B in 8 min; hold for 2 min and re-equilibrate to the initial conditions in 4 min. Starting condition of 40 % solvent B was utilized for long-chain bases (LCBs). For multiple reaction monitoring (MRM), precursor ions were [M+H]^+^ and product ions were dehydrated LCB fragments for ceramides and glycosylceramides. The loss of phosphoinositol-containing head groups was used for detection of glycosyl inositol phosphorylceramides.

#### Fatty acid methyl ester (FAME) derivatization and measurement by gas chromatography with flame ionization detection (GC-FID)

20 % of the total lipid extract was used for methylesterification to FAMEs, and total fatty acid quantification for normalizing the peak areas of individual lipid species measured by high performance UPLC-nanoESI-MS/MS. Aliquots were dried under nitrogen gas, and resuspended in a methanolic sulfuric acid solution. 5 µg tripentadecanoin standard was added to each sample, and the derivatization was carried out for 1 h in an 80 °C water bath. The reaction was stopped by addition of 200 µL saturated, aqueous NaCl, and the FAMEs extracted in hexane. The final extract was dried under nitrogen, and re-suspended in acetonitrile for injection into an Agilent 6890N GC-FID.

## Supplementary Data

**Supplementary Figure S1:** Sphingolipid content of mutant gametophores.

**Supplementary Figure S2:** Abnormal aniline blue stained accumulations occur in gametophores of the *s4h* mutant.

**Supplementary Figure S3:** Frequently-observed incomplete cell divisions in the *ipcst.92* mutant are identifiable in Z-stacks and fluorescence redistribution after photobleaching (FRAP) experiments.

**Supplementary Video SV1:** Representative fluorescence redistribution after photobleaching experiment of an *ipcst.92* phyllid cell. The photobleached region is outlined in red, and a nearby region used as reference for changes in carboxyfluorescein fluorescence over time without bleaching is outlined in blue.

## Acknowledgments

We thank Beate Preitz for support using the Zeiss LSM980 of the Developmental Biology Department of the University of Göttingen, and acknowledge the Imaging Unit of the JLU Giessen Germany for providing access to the TEM facilities. We thank Dr. Patricia Scholz for providing the R script used for statistical analysis, and Dr. Jasmin Gömann for the *s4h* mutant used in these experiments.

## Author Contributions

Author contributions: LW, CH, and TMH performed experiments; LW, CH, IF, KE, and TMH analyzed the data. TMH wrote the manuscript. LW, CH, IF, and KE edited the manuscript. IF, KE, and TMH designed and supervised the study. TMH agrees to serve as the author responsible for contact and ensures communication.

